# Curated Residual Decomposition for Increased MU Yield from HD-sEMG

**DOI:** 10.64898/2026.06.10.731304

**Authors:** Marius Osswald, Alessandro Del Vecchio

**Author notes:** The authors declare no competing financial interests.

## Abstract

High-density surface electromyography (HD-sEMG) decomposition algorithms identify motor units (MUs) from non-invasive recordings, yet even gold-standard methods typically decompose only a subset of all physiologically active MUs and preferentially extract higher-amplitude units, leaving low-amplitude MUs underrepresented. We propose Curated Residual Decomposition (CRD): Through decomposition of the residual signal remaining from a manually edited initial decomposition, we identify additional, mainly lower amplitude MUs that were previously undetected due to action potential superposition and preferential convergence to larger MUs.

We applied this approach to three datasets: concurrent HD-sEMG and HD-iEMG recordings from three muscles for two-source validation, and two publicly available datasets. Across all datasets, CRD substantially increased MU yield. In the two-source validation dataset, the first CRD iteration increased yield by 31–50% (FDI: 46.9%, TA: 49.8%, VL: 31.1%), with validation via intramuscular recordings confirming comparable identification accuracy of the CRD compared to the original MUs (RoA original: 0.994 ± 0.005; RoA CRD: 0.992 ± 0.009). In public datasets, yield increases ranged from 35–142% per muscle in the first iteration. Correspondingly, the explained signal power ratio increased, with the first iteration accounting for most gains. A second CRD iteration resulted in an additional, yet smaller yield increase (0–26.7%). Additionally identified MUs had smaller action potential amplitudes and lower recruitment thresholds, indicating the recovery of smaller, lower-threshold MUs.

CRD reliably increased MU yield and explained signal power across diverse muscles, datasets, and decomposition implementations. The approach provides an optional, validated step for more complete decomposition results across the full recruitment range in offline analyses.

## I. Introduction

Understanding the organization and control properties of the motor unit pool is fundamental to motor control research and clinical neuromuscular assessment. Motor units, which consist of a motoneuron and its muscle fibres, follow the Henneman size principle: smaller units activate first during weak contractions, and larger ones recruit as force increases [1]. The discharge patterns of individual motor units encode the motor commands issued by the nervous system and directly determine force production and control [2]–[4]. Consequently, access to representative samples of motor unit activity across the recruitment spectrum is essential for understanding motor control strategies, detecting neuromuscular pathology, and designing effective rehabilitation interventions.

High-density surface electromyography (HD-sEMG) has emerged as a non-invasive tool for recording motor unit activity across the entire muscle surface. By combining recordings from tens to hundreds of electrodes with blind-source-separation decomposition algorithms, such as convolutive kernel compensation (CKC) or Independent Component Analysis (ICA) variants, researchers can identify and track the discharge times of dozens to hundreds of individual motor units without the constraints and invasiveness of intramuscular recordings [5]–[9]. This capability has revolutionized the study of motor control, aging, and disease in human populations. However, the effectiveness of current decomposition approaches remains fundamentally limited to the observation of only a subset of all physiologically active MUs, with the proportion of the subset varying across experimental conditions, recorded muscle and participant [10], [11]. This holds true even for current state-of-the-art methods reporting particularly high MU yields [9], [12].

Blind-source-separation algorithms typically identify motor unit components in order of signal variance or dominance, and consequently preferentially extract high-amplitude motor units first. Motor units with smaller action potentials due to small motor neuron size, smaller muscle fibre territories, or location deeper within the muscle tissue are obscured by the superposition of larger action potentials from higher-threshold units, making them challenging to identify for decomposition algorithms, even when their activity is present in the original EMG signal [11]–[14]. This limits the number of MUs typically identified from HD-sEMG in submaximal contractions, especially at the lower end of the recruitment range, which is problematic as low-threshold MUs are highly relevant to low-force, daily motor tasks [1], [4], [15].

Approaches to reduce repeated convergence of decomposition algorithms on more dominant, larger sources through source deflation or iterative automated peel-off of the already identified spike trains have been proposed previously [7], [8], [16]–[18], but existing methods typically subtract uncurated (prior to manual editing) motor unit spike trains from the signal between iterations of the automatic decomposition algorithm. Because unedited spike trains contain errors such as spurious spikes, mixed sources and misidentified or missing discharges that have to be manually corrected after the automatic decomposition [11], [18], each successive peel-off step accumulates reconstruction error, progressively degrading the quality of the residual signal and ultimately limiting the yield improvement [8], [16], [19]. In contrast, if spike trains are manually curated to ensure accuracy before the residual signal is constructed, the residual signal should offer a much cleaner base for subsequent decomposition iterations. However, the effectiveness of such an iterative approach has not yet been performed or reported in scientific EMG literature.

In the present work, we propose the Curated Residual Decomposition (CRD) approach. We hypothesize that an iterative decomposition of the residual HD-sEMG signal reconstructed after manual edition of MU spike trains substantially increases motor unit yield while recovering physiologically valid, lower-threshold motor units across a range of experimental conditions and muscles. We test the approach using a two-source validation dataset of concurrent HD-sEMG and HD-iEMG recordings, which allows us to validate the additionally identified MUs against an independent reference, and two publicly available HD-sEMG decomposition datasets that span multiple muscles, participants, decomposition implementations, and manual editing operators. The aim is to establish CRD as a practical, validated and optional plug-in technique that can be integrated into any offline decomposition workflow for improving the completeness of motor unit decomposition from HD-sEMG.

## II. Methods

### A. Datasets

#### 1) Two-Source Validation Dataset

Concurrent HD-sEMG and HD-iEMG data from three different muscles was acquired (Fig. 1B). 16-channel HD-iEMG electrode arrays designed for acute recordings [8], [20] were implanted into the muscle by a medical professional under ultrasound guidance. HD-iEMG, HD-sEMG and force signals were recorded using a multi-channel amplifier (Quattrocento, OT Bioelettronica, Turin, Italy) at a sampling rate of 10,240 Hz with 16-bit resolution in a monopolar configuration. All data acquisitions were approved by the ethics committee of the Friedrich–Alexander University Erlangen–Nürnberg (approval number 23-271-B) and were conducted in accordance with the Declaration of Helsinki.

**Fig. 1.**
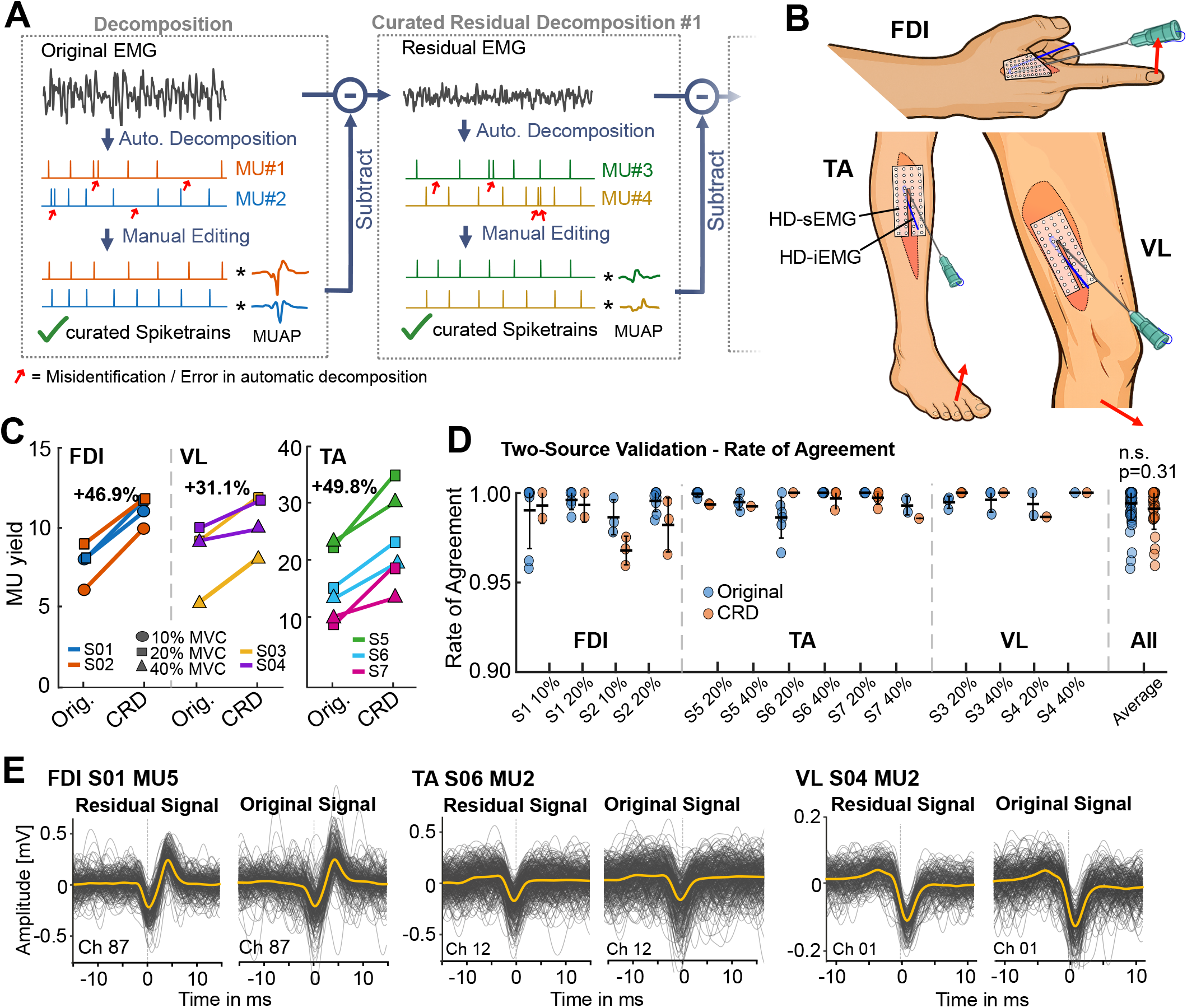
Curated Residual Decomposition Pipeline and Validation. A: Schematic process of the Curated Residual Decomposition approach. Original decomposition results are manually edited to obtain valid MU spike trains. The spike trains are convolved with their MUAPs and subtracted from the original EMG signal to receive the residual EMG unexplained by the original decomposition. This residual signal is then decomposed and after manual editing additional MUs extend the original decomposition’s MU pool. This process can be repeated for further increased MU yield. B: Experimental setup of HD-sEMG and HD-iEMG acquisition for two-source validation in FDI, TA and VL. C: Identified MUs from the HD-sEMG data in the two-source validation dataset split into the three muscles, separately for each recording (left), and summarized into relative MU yield increase per muscle (right). D: The two-source validation shows similar rate of agreements between the intramuscular MUs and the MUs obtained from the original sEMG decomposition (blue) and the sEMG MUs obtained from CRD (orange). E: Shimmer plots of sEMG MUs obtained from CRD using the residual EMG signal vs. the original EMG signal. Plots show the MUs have been present in the original EMG already, but under increased superpositions of other MU action potentials.

##### First Dorsal Interosseus

Two participants (28 and 29 years, 1 male, 1 female) performed isometric index finger abduction at 10 and 20% of their maximum voluntary contraction (MVC) for 20 s while fixed in a custom-built dynamometer for index finger contractions [21]. The HD-iEMG arrays were implanted in the First Dorsal Interosseus (FDI) in an oblique direction from the distal third of the muscle towards its proximal third (see [22] for details), roughly parallel to the skin surface covered by the HD-sEMG grid. HD-sEMG was recorded using 96ch electrode grids with an IED of 3 mm, specifically designed to cover the entire FDI surface (GR03MM1807, OT Bioelettronica, Turin, Italy). The reference electrode was placed around the participant’s wrist.

##### Tibialis Anterior

Three participants (mean age 28.6 years, 3 male) performed isometric ankle dorsiflexion at 20 and 40% of MVC for 20 s while fixed in a foot dynamometer. The HD-iEMG electrode arrays were implanted in muscle belly of the Tibialis anterior (TA) at an approximate angle of 45° relative to the skin surface. HD-sEMG was recorded with a 64ch electrode grid with an IED of 8mm (GR08MM1305, OT Bioelettronica, Turin, Italy). To maximize the chance of MU matches between the surface and intramuscular signals, a slit was cut in the center of the sEMG grids which served as the implantation site of the intramuscular arrays. The reference electrode was placed around the ankle joint.

##### Vastus Lateralis

Two participants (33 and 33 years, 2 male) performed isometric leg extension at 20 and 40% MVC for 20 s while fixed in a custom built leg dynamometer at a knee flexion angle of 45°. HD-sEMG was recorded from the Vastus Lateralis (VL) with 64ch electrode grids with an IED of 8mm (GR08MM1305, OT Bioelettronica, Turin, Italy). The reference electrode was placed around the ankle joint. Analogous to the TA data, the HD-iEMG electrode arrays were implanted in muscle belly at an approximate angle of 45° relative to the skin surface with a slit in the center of the HD-sEMG grid serving as the implantation site of the intramuscular arrays.

#### 2) Public HD-sEMG Decomposition Datasets

To check the efficiency of the proposed CRD approach on a wider variety of muscles and experimental conditions, a randomly selected subset of two publicly available HD-sEMG datasets was used. For detailed descriptions of the electrode placement, data acquisition and experimental protocols, we refer to the original publications.

##### Avrillon et al. (2024)

Data from Avrillon et al. [9] were used, publicly available under a CC BY 4.0 license [23]. The dataset contains HD-sEMG recordings from the tibialis anterior and vastus lateralis muscles during isometric contractions at various force levels up to 80% of maximum voluntary contraction (MVC). Monopolar HD-sEMG was recorded with four 64ch electrode grids in a 2 × 2 arrangement with 4mm IED for the TA and 8mm for the VL. The authors provide manually edited decomposition results of all recordings. This dataset shows an exceptionally high average MU yield in both recorded muscles with 129 ± 44 MUs per participant in the TA and 130 ± 63 in the VL [9]. For the present study, the published edited decomposition results served as the starting point for the CRD pipeline described below, and the same MU quality thresholds were applied as in the original publication. We used TA data at 20, 40 and 60% of MVC from four randomly selected participants (S1, S2, S4 and S6) and VL data at the same contraction intensities from three randomly selected participants (S11, S13, S17). As the authors did not publish participant-specific biometric information, we cannot provide sex or mean age of this subset of participants and refer to the overall biometrics of the entire dataset as published in [9].

##### Hug et al. (2021)

Data from Hug et al. [24] were used, publicly available under a CC BY 4.0 license [25]. The dataset contains HD-sEMG recordings from three lower-leg muscles: gastrocnemius medialis (GM), gastrocnemius lateralis (GL) and TA. For all muscles, monopolar HD-sEMG was recorded with one 64ch electrode grid with an IED of 8 mm (ELSCH064NM2, SpesMedica, Battipaglia, Italy). The authors provide the manually edited decomposition result from 8 trained experts editing the decompositions independently. In this study, we included the data of operator 8, as this operator reported the most experience with the process. In addition to the manually edited decomposition results, the authors also provide the unedited automatic decomposition results, which allowed us to further validate the novelty of the additional MUs identified during the CRD. During the editing of the residual decompositions, the same MU quality thresholds were applied as reported by Hug et al. [24]. Biometric details of the dataset can be found in the original publication.

### B. Signal Processing and Decomposition

#### 1) MU Decomposition and Manual Editing

The initial decomposition of the two-source validation dataset, as well as the CRD iterations of all three datasets, were performed using MUedit [18], an implementation of a convolutive blind source separation approach for MU identification from HD-iEMG and HD-sEMG signals [8]. Automatic peel-off of identified MUs after each decomposition iteration was applied, an initial silhouette value threshold of 0.88 set and duplicate MUs automatically removed after completed automatic decomposition. Briefly, the algorithm extends the multichannel surface EMG observation by convolutive whitening and iteratively estimates a set of linear separation filters that maximise the sparseness of the projected signal, thereby identifying the discharge times of individual MUs.

Following the automatic decompositions, the resulting MU pulse trains were manually inspected and edited using the DEMUSE toolbox to correct instances of misidentified MU firings. This process is necessary even in signals of high signal quality and is described in detail in Del Vecchio et al. [11].

Different decomposition quality metrics were applied for the exclusion of MUs for further analysis and signal deflation were applied, depending on the approaches reported in the two public datasets. Hug et al. [24] use a pulse-to-noise ratio of 30 dB as threshold, which therefore was applied here to this dataset’s CRD iterations, as well as to all decompositions of the two-source validation dataset. Avrillon et al. [9] use a post-editing silhouette value of above 0.9 as quality threshold, which was hence similarly applied to the edited residual decompositions of this dataset in this study.

#### 2) Curated Residual Decomposition

In brief, the here presented CRD approach subtracts manually edited MU spike trains convolved with their MUAP templates from the original EMG signal, decomposes the resulting residual signal followed by manual editing of the result spike trains to receive additional MUs to add to the identified MU pool (see Fig. 1A).

For each originally decomposed MU that passed the quality threshold, a motor unit action potential (MUAP) template was estimated via spike-triggered averaging (STA). A window of ±12.5 ms was centred on each discharge time, and the waveforms across all active electrode channels were averaged over valid discharge events (discharges within 25 ms of the signal edges were excluded from averaging). Templates were estimated and subtracted sequentially: for MU *k*, the STA was computed from the signal after the contributions of all preceding MUs had already been removed, and the resulting template was then subtracted at every discharge time of MU *k* (peel-off). Performing this procedure for all *N* MUs yields the residual signal, which represents the HD-sEMG activity not accounted for by the identified MU pool of the initial decomposition and all potential previous CRD iterations. This process is similar to the processing step often referred to as peel-off, where identified MUs are removed from the signal after each iteration of the automatic decomposition algorithm to avoid the repeated convergence onto the same sources in subsequent iterations [7], [12], [16]–[18]. However, in our approach we ensure only valid MU spike trains are subtracted, that have been edited and curated by a trained expert and therefore do not induce cumulative errors in the remaining signal due to misalignment of action potentials or the subtraction of misidentified firings or mixed sources [8], [18].

The residual signal was decomposed using the same implementation and parameters described above, followed by the same manual editing and PNR quality-control procedure. To exclude duplicate MUs, each newly identified MU was compared against all previously identified MUs of the initial and all potential previous CRD iterations by computing the maximum normalised cross-correlation of binary spike trains within a ±200 ms lag window. MU pairs with a value above 0.20 were considered duplicate.

For the two-source validation dataset, CRD was performed once. For the two public datasets [9], [24], a second CRD iteration was performed.

#### 3) Two-Source Validation of CRD MUs

To assess the validity of MUs identified through the CRD, concurrently recorded HD-iEMG signals were used as an independent reference for the MUs detected from HD-sEMG. Surface and intramuscular MU spike trains were first matched using normalised cross-correlation over a lag range of ±0.5 s; a pair was accepted as originating from a common MU if the peak cross-correlation exceeded 0.2. This threshold would allow MU pairs to be included in the subsequent rate of agreement (RoA) analysis, even if only a subset of their firings would match, as it would be the case for poorly identified MUs, or two separate MUs being identified as one (mixed sources). For matched pairs, the RoA was computed as

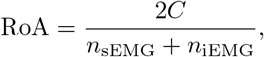

where *C* is the number of coincident spikes within a ±0.5 ms tolerance window and *n* the total number of spikes in the respective sEMG and iEMG spike trains [26]. All RoA values were computed within the constant-force plateau phase of the contractions. RoA distributions were compared between MUs from the original decomposition and those identified in the CRD to determine whether the additional MUs achieved comparable matching quality.

#### 4) Proportion of Explained Signal Power Analysis

To quantify the cumulative contribution of the identified MU pool to the recorded HD-sEMG signal, we computed the explained signal power ratio for each MU set (original, CRD 1 & CRD 2), following the approach described in Avrillon et al. [9].

For a given MU set, the signal was reconstructed by placing each MU’s STA template at every discharge time and summing the contributions of all MUs. The explained signal power ratio was defined as the mean squared reconstructed signal *x*_resynth_ relative to the mean squared original signal *x*:

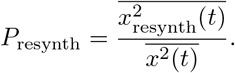

The ratios were averaged across all active electrode channels and computed within the constant-force plateau epoch to avoid contributions from force ramps.

#### 5) CRD MUs’ Depth and Size

As we hypothesized the MUs identified in the residual decompositions to show smaller signal amplitudes due to either being located deeper in the muscle or of smaller MU size, we compared the MUAP amplitude and recruitment thresholds of all MUs of the original and combined CRD iterations of all three datasets. As the MUAPs are low-pass filtered and attenuated on the way from the muscle fiber to the electrode sites, deeper MUs will naturally show smaller MUAP amplitudes, given similar MU sizes [2]. Conversely, smaller MUs typically show smaller MUAPs due to a lower number of innervated muscle fibers, as well as lower recruitment thresholds as dictated by Henneman’s size principle [1]. MUAP amplitude was quantified as the mean peak-to-peak amplitude of the spike-triggered average template averaged across the three largest amplitude channels. Recruitment threshold was defined as the force level in % MVC at which the motor unit discharged for the first time within the recorded contraction. As a second estimate of MU depth, in the two-source validation dataset we checked the channel in the linear 16-channel HD-iEMG array where the MUs showed their largest MUAP amplitude. Deeper MUs should have their center of activity, i.e. largest MUAP amplitude, in more distal array channels. This analysis was not performed for the FDI data, as the intramuscular arrays there were implanted approximately in parallel to the skin surface, contrary to the approx. 45° implantation angle for the TA and VL muscles.

### C. Statistical Analysis

MUAP amplitude and recruitment threshold were compared across original decomposition vs. CRD iterations using linear mixed models (LMM). Decomposition group (Original, CRD) was entered as a categorical fixed effect with the original decomposition as the reference level. Recording and participant were included as random intercepts to account for the non-independence of MUs recorded within the same recording or participant (value ∼ group + (1 | participant) + (1 | recording)). The significance of the Group fixed effect was assessed by an F-test. Pairwise differences between groups were evaluated using contrast matrices, and p-values were Bonferroni-corrected for the number of comparisons. Analyses were performed separately for each muscle and dataset.

To compare the depth of original versus residual MUs within the intramuscular array, the dominant channel of the 16-electrode linear intramuscular EMG array (channel 1 = deepest; channel 16 = shallowest) was identified for each matched MU as the channel exhibiting the largest spike-triggered average peak-to-peak amplitude. Statistical comparison of dominant channel depth between original decomposition and CRD iterations was performed using the Wilcoxon signed-rank test. To avoid pseudo-replication arising from multiple MUs contributing to a single contraction, the median dominant channel was computed per recording for each decomposition group, and the signed-rank test was applied to these per-recording medians (one observation per recording).

Raw p-values were Bonferroni-corrected and significance was declared at *α* = 0.05 (* p *<* 0.05, ** p *<* 0.01, *** p *<* 0.001).

## III. Results

### A. Two-Source Validation

The Curated Residual Decomposition increased the number of identified HD-sEMG MUs in all muscles of the two-source validation dataset (Fig. 1C, Table I). As commonly observed in HD-sEMG decomposition [11], [12], [24], MU yield varied across muscles (original decomposition yield per recording: TA = 9–23 MUs, VL = 5–10 MUs, FDI = 6–9 MUs, see Fig. 1 and Table I). One iteration of the approach increased the number of identified MUs by approximately 30–50% relative to the original decomposition (FDI: 46.9%, TA: 49.8%, VL: 31.1%), corresponding to 3–4 additional MUs in the FDI, 3–12 in the TA, and 1–3 in the VL (Table I).

**TABLE 1.** Two-source Validation Dataset. MU yield and rate of agreement to HD-iemg mus of MUs found in the original HD-semg decomposition and CRD. True new CRD MUs are MUs identified in the residual decomposition that were not present in the original decomposition prior to manual editing.

| Muscle | Part. | Contr. Level | # intram. MUs | # surface MUs original | # common MUs original | mean RoA original | # surface MUs CRD | # common MUs CRD | mean RoA CRD | # true new CRD MUs |
| --- | --- | --- | --- | --- | --- | --- | --- | --- | --- | --- |
| FDI | S1 | 10 | 19 | 8 | 7 | 0.988 | 3 | 2 | 0.993 | 2 |
|  |  | 20 | 27 | 8 | 6 | 0.996 | 4 | 2 | 0.993 | 4 |
|  | S2 | 10 | 29 | 6 | 3 | 0.986 | 4 | 3 | 0.967 | 1 |
|  |  | 20 | 28 | 9 | 8 | 0.995 | 3 | 3 | 0.982 | 0 |
| TA | S5 | 20 | 21 | 22 | 6 | 0.999 | 12 | 2 | 0.993 | 12 |
|  |  | 40 | 24 | 23 | 4 | 0.995 | 7 | 1 | 0.992 | 6 |
|  | S6 | 20 | 28 | 16 | 6 | 0.986 | 6 | 1 | 1.000 | 2 |
|  |  | 40 | 31 | 13 | 5 | 1.000 | 6 | 3 | 0.997 | 3 |
|  | S7 | 20 | 31 | 9 | 3 | 1.000 | 9 | 5 | 0.997 | 6 |
|  |  | 40 | 34 | 10 | 2 | 0.993 | 3 | 1 | 0.985 | 1 |
| VL | S3 | 20 | 6 | 10 | 3 | 0.994 | 2 | 2 | 1.000 | 2 |
|  |  | 40 | 5 | 9 | 3 | 0.996 | 1 | 1 | 1.000 | 0 |
|  | S4 | 20 | 10 | 9 | 2 | 0.994 | 3 | 1 | 0.986 | 1 |
|  |  | 40 | 14 | 5 | 1 | 1.000 | 3 | 1 | 1.000 | 3 |
| Mean ± SD |  |  |  |  |  | 0.994 ± 0.005 |  |  | 0.992 ± 0.009 |  |

To validate that the additional MUs were correctly identified, their RoA against the intramuscular reference was compared against the original decompositions. On average across all recordings, 4.2 ± 2.1 MUs from the original decomposition and 2.0 ± 1.2 MUs from CRD were also identified in the intramuscular reference (Table I). RoA values were high for both sets: 0.994 ± 0.005 for originally decomposed MUs and 0.992 ± 0.009 for MUs from CRD (Fig. 1D). The difference was not statistically significant (Wilcoxon signed-rank test: *p* = 0.31, Cohen’s *d* = 0.31), indicating that CRD identified MUs with discharge-timing accuracy comparable to that of the original decomposition.

Shimmer plots further confirmed that the MUs recovered by CRD were genuinely present in the original HD-sEMG signal already, not merely artefacts of the residual computation. When STA templates of residual MUs were aligned to their discharge times in the original signal, clear MUAP waveforms were visible across the electrode grid (Fig. 1E, right column), albeit with reduced signal-to-noise ratio compared to the residual signal from which they were identified (Fig. 1E, left column).

To confirm that CRD identified genuinely novel MUs rather than MUs previously rejected during manual editing due to insufficient PNR, the MUs identified in CRD were matched against the unedited automatic output of the original decomposition, using the same normalized-cross-correlation approach as described above. This was performed for the two-source validation and the Hug (2021) datasets. Avrillon (2024) did not provide unedited decomposition results. The number of true new MUs in each datafile is reported in Table I for the two-source validation dataset and Table II for Hug (2021). In most recordings, the majority of MUs identified during CRD iterations were not part of the original unedited decomposition result.

**TABLE 2.** Public datasets Avrillon (2024) And Hug (2021). MU yield in the original HD-sEMG decomposition and CRD. True new CRD MUs in Hug (2021) Are CRD it. 1 & 2 MUs not present in the original decomposition prior to manual editing.

**Dataset Avrillon (2024)**
| Muscle | Part. | Contr. Level | # MUs original | # MUs after CRD It.1 | # MUs after CRD It.2 |
| --- | --- | --- | --- | --- | --- |
| TA | S1 | 20 | 80 | 106 (+26) | 112 (+6) |
|  |  | 40 | 83 | 106 (+23) | 114 (+8) |
|  |  | 60 | 77 | 86 (+9) | 96 (+10) |
|  | S2 | 20 | 83 | 112 (+29) | 116 (+4) |
|  |  | 40 | 93 | 126 (+33) | 132 (+6) |
|  |  | 60 | 80 | 101 (+21) | 110 (+9) |
|  | S4 | 20 | 48 | 65 (+17) | 72 (+7) |
|  |  | 40 | 51 | 75 (+24) | 82 (+7) |
|  |  | 60 | 28 | 51 (+23) | 58 (+7) |
|  | S6 | 20 | 30 | 38 (+8) | 40 (+2) |
|  |  | 40 | 27 | 36 (+9) | 37 (+1) |
|  |  | 60 | 21 | 27 (+6) | 27 (+0) |
| VL | S11 | 20 | 44 | 72 (+28) | 80 (+8) |
|  |  | 40 | 48 | 86 (+36) | 97 (+11) |
|  |  | 60 | 28 | 42 (+14) | 48 (+6) |
|  | S13 | 20 | 60 | 109 (+49) | 124 (+15) |
|  |  | 40 | 46 | 70 (+24) | 81 (+11) |
|  |  | 60 | 38 | 65 (+27) | 74 (+9) |
|  | S17 | 20 | 37 | 59 (+22) | 68 (+9) |
|  |  | 40 | 38 | 69 (+31) | 77 (+8) |
|  |  | 60 | 34 | 38 (+4) | 41 (+3) |

**Dataset Hug (2021)**
| Muscle | Contr. Level | # MUs original | # MUs after CRD It.1 | # MUs after CRD It.2 | # true new CRD MUs |
| --- | --- | --- | --- | --- | --- |
| GL | 10 | 4 | 5 (+1) | 5 (+0) | 0 |
|  | 30 | 9 | 13 (+4) | 13 (+0) | 1 |
|  | 50 | 4 | 8 (+4) | 8 (+0) | 1 |
|  | 70 | 5 | 5 (+0) | 5 (+0) | 0 |
| GM | 10 | 17 | 30 (+13) | 32 (+2) | 11 |
|  | 30 | 27 | 60 (+33) | 72 (+12) | 42 |
|  | 50 | 15 | 38 (+23) | 39 (+1) | 24 |
|  | 70 | 6 | 15 (+9) | 16 (+1) | 7 |
| TA | 35 | 21 | 52 (+31) | 56 (+4) | 35 |
|  | 50 | 15 | 33 (+18) | 40 (+7) | 25 |
|  | 70 | 7 | 18 (+11) | 19 (+1) | 12 |

### B. Effectiveness of Curated Residual Decomposition

CRD consistently increased the number of identified MUs across both public datasets and all recorded muscles. The magnitude of the yield increase was substantial after the first CRD iteration, whereas the second iteration provided a smaller but still measurable additional gain (Fig. 2A–C, Table II).

**Fig. 2.**
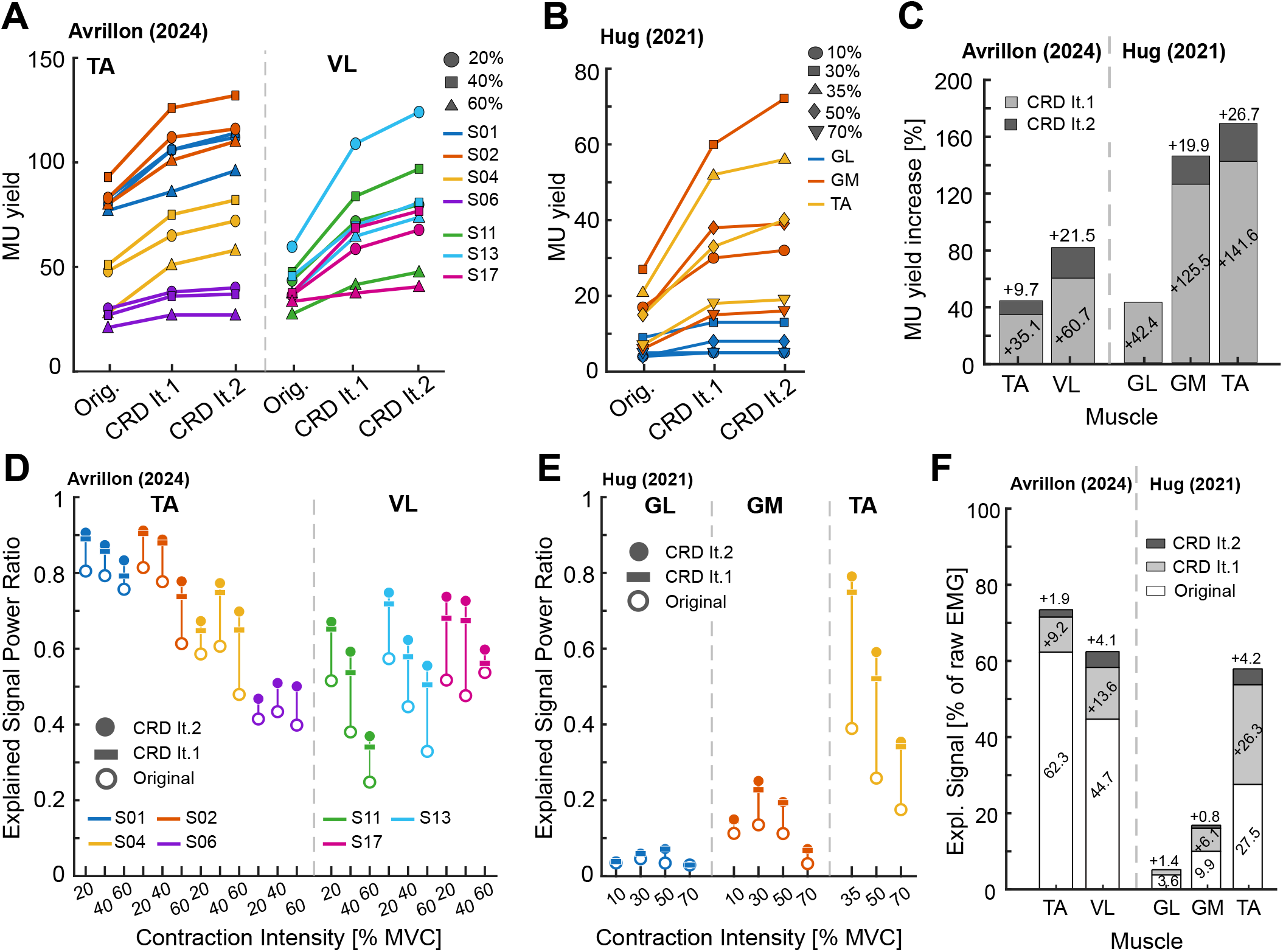
Increased MU yield and explained signal. A: From dataset Avrillon (2024): Absolute number of decomposed MUs in TA (left) and VL (middle) after original decomposition and first and second CRD. B: Analogue to panel A for dataset Hug (2021). C: Increase of MU yield after first and second CRD iteration relative to the original decomposition yield for both datasets, split into muscles. D: Ratio of the signal’s power explained by the sum of all identified MUs to the original EMG signal’s power for the data from Avrillon (2024). E: Analogue to panel D for Hug (2021). F: Average ratio of explained signal power in percentage after original decomposition, and increased percentage of explained signal after CRD, for both datasets split into muscles.

In the Avrillon (2024) dataset, the publicly available original decomposition identified on average 58.4 ± 26.9 MUs per recording in the TA and 41.4 ± 9.3 MUs in the vastus lateralis (VL) across participants. Across all subjects and contraction intensities, the first iteration of CRD increased the originally published MU yield by 35.1 ± 17.0% in the TA and 60.7 ± 21.7% in the VL (additional 6–33 TA, 4–49 VL MUs). A second CRD iteration increased the yield by another 9.7 ± 6.5% for the TA, and 21.0 ± 5.0% for the VL, relative to the original decomposition yield (additional 0–10 TA, 3–15 VL MUs, see Fig. 2A).

The data provided by Hug (2021) had a lower initial decomposition yield compared to Avrillon (2024) with an average initial MU yield of 5.5 ± 2.4 for the GL, 16.3 ± 8.6 in the GM and 14.3±7.0 in the TA. CRD iteration 1 increased the MU yield by 42.4±42.4% in the GL, 125.5±35.5% in the GM and 141.6 ± 19.3% in the TA, as well as 19.9 ± 16.9% (GM) and 26.7 ± 17.5% (TA) after the second iteration. Iteration 2 did not identify additional MUs that surpassed the MU quality threshold for the GL (Fig. 2B).

Generally, across all datasets and muscles, the number of additionally identified MUs increased relative to the initial yield from the original decomposition (see Fig. 2A and B). As an example, in Avrillon (2024) TA subject S6 with an average of 26.0 ± 4.6 MUs originally decomposed, both CRD iterations identified a combined additional number of 8.7 ± 2.3 MUs, while S2 with the much higher number of 85.3 ± 6.8 initial MUs led to 34.0 ± 4.6 additional MUs after both CRD iterations.

The increase in MU yield was accompanied by a corresponding increase in the proportion of original signal power explained by the identified MUs (Fig. 2D–F). In the Avrillon (2024) dataset, the explained signal ratio after the original decomposition was 62.3±16.3% in the TA and 44.7±10.8% in the VL (mean ± SD across recordings; Fig. 2D). The first CRD iteration increased the explained ratio by 9.2 ± 3.9 percentage points (pp) in the TA and 13.6 ± 5.1 pp in the VL. The second iteration yielded a further modest increase of 1.9±1.7 pp (TA) and 4.1 ± 1.4 pp (VL; Fig. 2D and F).

An analogous pattern was observed in the Hug (2021) dataset (Fig. 2E and F): the explained ratio after the original decomposition was 3.6 ± 0.7% (GL), 9.9 ± 4.5% (GM), and 27.5 ± 10.8% (TA). The first CRD iteration increased the ratio by 1.4 ± 1.7 pp (GL), 6.1 ± 3.1 pp (GM), and 26.3 ± 9.7 pp (TA); the second iteration added 0.8 ± 1.0 pp (GM) and 4.2 ± 2.9 pp (TA).

Across both datasets, the first CRD iteration accounted for the large majority of the total explained signal gain, while the second iteration contributed a consistently smaller increment (Fig. 2F). Generally, muscles and recordings with a larger relative increase in MU yield also showed a larger increase in explained signal power, suggesting that the additional MUs identified through CRD make a proportionate contribution to the recorded EMG (Fig. 2C and F).

### C. Curated Residual Decomposition Yields Smaller, Lower-Threshold MUs

To check if CRD facilitates the identification of MUs of a certain size, we compared the MUAP amplitude and recruitment threshold of the MUs found by the original decompositions vs. the CRD iterations across all datasets and muscles.

Figures 3A–B visualize the result on a MU level as well as per-recording level. Across all datasets and muscles, both the median MUAP amplitudes and recruitment thresholds of the curated residual CRD were smaller than for the MUs of the original decompositions. Statistical significance was reached for the MUAP amplitude of the FDI in the two-source validation dataset (*β* = −65.4 *μ*V, 95% CI [−89.8, −41.1], *t*(134) = −5.3, *p <* 0.001) and the GM and TA of the Hug (2021) dataset (GM: *β* = −285.5 *μ*V, 95% CI [−360.2, −210.7], *t*(157) = −7.49, *p <* 0.001; TA: *β* = −114.9 *μ*V, 95% CI [−152.1, − 77.6], *t*(113) = −6.04, *p <* 0.001), as well as for the recruitment threshold of the FDI and TA in the two-source validation dataset (FDI: *β* = −1.16 %MVC, 95% CI [−1.91, −0.41], *t*(43) = −3.03, *p* = 0.024; TA: *β* = −3.34 %MVC, 95% CI [−5.50, −1.17], *t*(134) = −3.01, *p* = 0.018), the TA in the Avrillon (2024) data (*β* = −8.51 %MVC, 95% CI [−10.15, −6.87], *t*(994) = −10.18, *p <* 0.001) and the GM and TA from Hug (2021) (GM: *β* = −5.71 %MVC, 95% CI [−8.92, −2.49], *t*(157) = −3.48, *p* = 0.004; TA: *β* = −11.18 %MVC, 95% CI [−15.00, −7.37], *t*(113) = −5.75, *p <* 0.001). A notable exception from the trend was the recruitment thresholds of the VL MUs at 40% MVC in the Avrillon (2024) data, which showed a slight increase in recruitment threshold in the CRD MUs across all three participants.

**Fig. 3.**
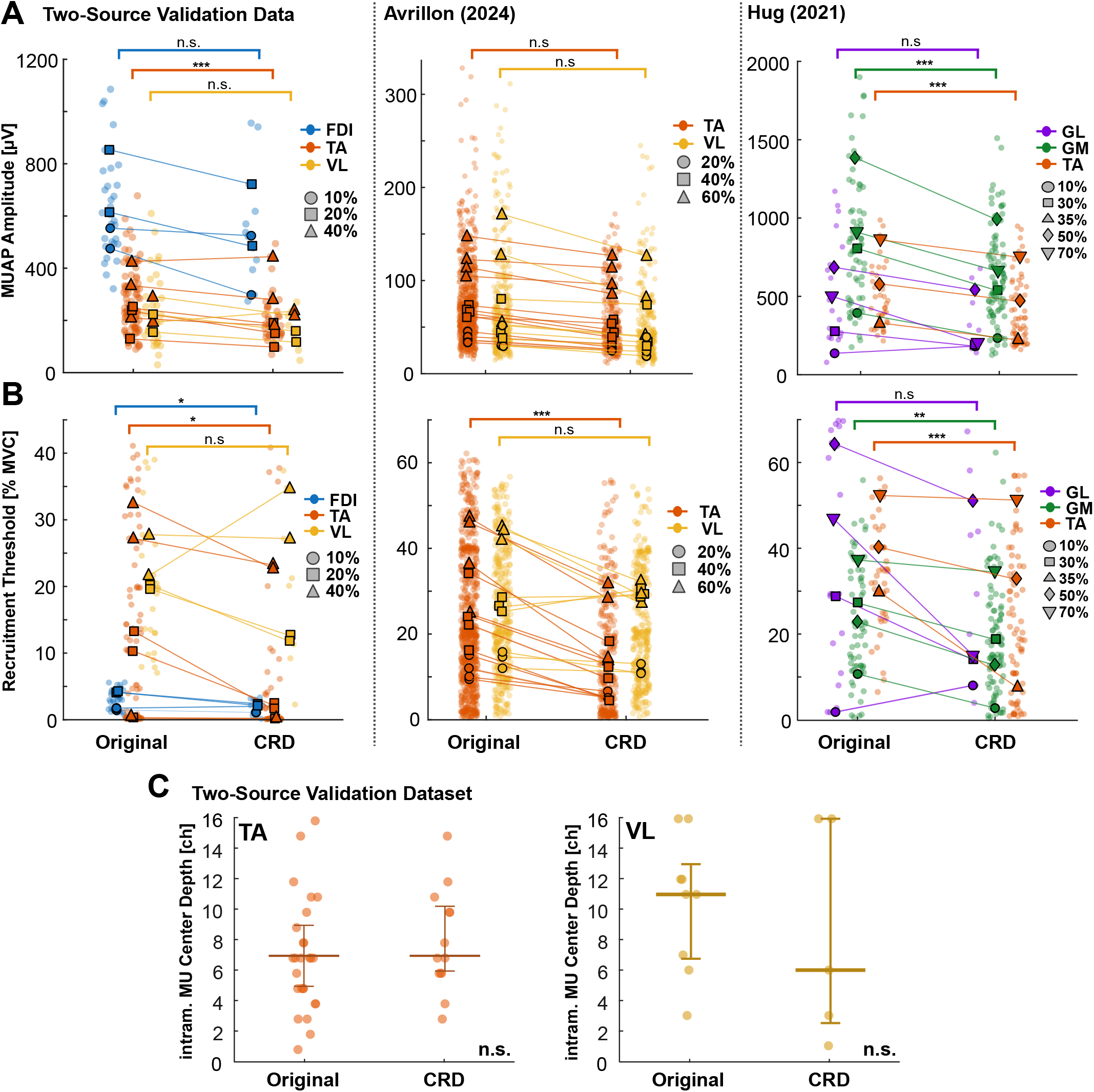
Size and depth of MUs from Curated Residual Decompositions. A: MUAP amplitudes of MUs from original decompositions (blue) and CRD iterations (red) of all three datasets. Dots mark single MUs, lines indicate the median MUAP amplitude of each recording after original decomposition and CRD. MUs identified in the CRD iterations show a consistently decreased MUAP amplitude, although the effect is not statistically significant in all muscles. Colors denote muscles. B: Analogue to A, but showing the recruitment thresholds of the identified MUs. A generally consistent decrease in recruitment threshold is observed. C: Data from the two-source validation dataset, showing the center of the MUs matched between sEMG and iEMG decompositions split into original sEMG decompositions and CRD. Channel index goes from highest to lowest, meaning ch16 is closest to the skin surface and ch1 deepest inside the muscle. MUs found in CRD do not show significantly increased depth within the muscle.

As the observed average reduction in MUAP amplitude could not only be explained by a generally smaller MU size, but also further muscle fiber-to-electrode distance, meaning deeper MU location within the muscle, we checked the depth of the MU centers-of-activation within the original decomposition and CRD MUs of the two-source validation dataset by comparing their location of largest MUAP channel within the intramuscular EMG electrode array. Only MUs matched between the sEMG and iEMG data were included. For the FDI data this analysis was not performed, as the intramuscular electrode arrays were placed mostly parallel to the skin surface. Interestingly, for both the TA and VL data, no significant effect of decomposition group on MU depth could be observed (Fig. 3C; TA: median original = 7.00, median redec. = 7.25, *W* = 6, *p >* 0.05; VL: median original = 11.50, median redec. = 7.75, *W* = 6, *p >* 0.05).

## IV. Discussion

Even current state-of-the-art HD-sEMG decomposition approaches that report particularly high MU yields [9], [12] explain only a fraction of the total recorded signal power and systematically underrepresent lower-amplitude MUs, as the dominant, high-amplitude sources drive convergence and mask the weaker contributions of smaller units. This is an inherent consequence of the signal superposition problem: once the high-amplitude MU pool has been identified, what remains in the signal is a complex mixture of low-amplitude activity that any single decomposition pass struggles to resolve. In this work, we demonstrate that iterative Curated Residual Decomposition substantially increases the number of identified MUs beyond what is recovered from a single decomposition pass, while also explaining a larger proportion of the recorded signal power. Two-source validation against concurrent HD-iEMG, a gold-standard for MU decomposition validation [8], [9], [12], [26]–[28], confirmed that the additional MUs are physiologically valid, with RoA values indistinguishable from those of originally decomposed MUs. Shimmer plots demonstrate that the additional MUs were already present in the original EMG signal but obscured by the superposition of higher-amplitude action potentials. Duplicate matching further ensured no redundant identification. The effect was consistent across three independent datasets including five lower- and upper-limb muscles, multiple contraction intensities, two distinct decomposition algorithm implementations, and several independent manual editing operators.

Iterative removal of identified sources from the EMG signal during MU decomposition is not a new idea [7], [12], [17], [18]. However, existing automatic peel-off pipelines subtract uncurated (pre manual editing) MU pulse trains from the signal between automatic decomposition iterations. Because misidentified spikes and missed discharges in the unedited pulse trains propagate into the residual signal after subtraction, each subsequent peel-off step accumulates an increasing reconstruction error, which is an issue previously acknowledged with different strategies proposed to overcome the limitation [8], [16]. The key methodological difference in the present work is that the spike trains are manually curated before the residual is constructed, so the residual signal is, to the best of the expert manual editing operator’s ability, the EMG signal minus the activity of a set of MUs whose discharge times are correctly identified. This residual signal therefore offers a cleaner substrate for a second decomposition pass, which we argue is the principal reason the approach consistently uncovers additional valid MUs. However, the present approach should not be considered a replacement of automatic peel-off but rather a complement. During the automatic decomposition of the original signals (two-source validation dataset) and curated residuals (all datasets) in this work, automatic peel-off was also performed, as the previous work cited above demonstrated its effectivity despite its limitations. We also acknowledge the effectiveness of the CRD approach may depend on the experience of the operator in the manual editing process, as errors in the edited spike trains will inevitably induce irreparable noise in the residual signal. Still, Hug et al. (2021) showed that given a certain level of experience, interoperator reliability in the manual edition is high.

Within our data, the first CRD iteration accounted for the clear majority of the total additional yield and explained-signal gain, while a second iteration produced a smaller but still measurable contribution. This pattern is expected: as successive iterations remove the dominant contributors to the residual, the signal-to-noise ratio available for separating remaining MUs progressively deteriorates. A third or fourth iteration would presumably provide further diminishing returns until the remaining signal is dominated by physiological noise, electrode noise, and crosstalk that cannot be attributed to a discrete MU. In practice, additional CRD iterations are therefore a less and less beneficial trade-off between yield gain and analyst effort.

The absolute yield gain was largest in recordings that already provided a medium to high original yield, and smaller where the original decomposition yielded only a small number of MUs (Fig. 2A and B, Table II, see Avrillon (2024) S1/S2 vs. S6, Hug (2021) GL vs. GM). This is intuitive: when the original decomposition identifies few MUs, the residual signal is dominated by unattributed MU activity that the same separation algorithm presumably could not isolate on the first pass either. Iterating the same algorithm does not, in itself, overcome the underlying recording-quality or anatomical limitations. Conversely, recordings with rich initial MU identification yield correspondingly larger absolute and relative gains because the deflation step more cleanly exposes the remaining minority sources. CRD thus complements, rather than replaces, efforts to improve the initial decomposition through better recording hardware, electrode placement, or algorithmic advances.

The additionally identified MUs were characterized by systematically smaller MUAP amplitudes and lower recruitment thresholds than the MUs from the original decomposition. These two characteristics are consistent with one another considering the size principle [1]. Lower-threshold motoneurons innervate smaller numbers of muscle fibres, producing smaller extracellular potentials at the skin surface [2]. Combined, the two observations support the interpretation that CRD preferentially recovers physiologically smaller MUs that are masked at the surface by the much larger action potentials of the high-threshold population. This finding is of direct physiological relevance for several reasons. The low-threshold MU pool is a critical determinant of fine force control and is typically underrepresented in HD-sEMG decompositions of submaximal contractions [9], [11], [29]. Because the effective neural drive to a muscle is the common synaptic input shared across the active motoneuron pool and is recovered with increasing fidelity as more of that pool is sampled, recovering a larger fraction of MUs improves the reconstruction of the effective neural drive [3], [12], [30].

The recovered units also extend the observable rate-coding spectrum as the firing behaviour of early-recruited MUs differs qualitatively from that of later-recruited units [9], [31], [32], so an unbiased view of rate coding of the full MU pool requires sampling both ends of the recruitment range. Finally, there is evidence neuromuscular pathology and ageing preferentially affect low-threshold MUs [33], [34], so a more complete sample of this MU sub-population is particularly valuable in clinical and ageing-research contexts.

A plausible alternative explanation for reduced MUAP amplitude is increased fibre-to-electrode distance, since the soft tissue between the muscle and the skin, as well as the muscle itself act as a spatial low-pass filter that attenuates deeper sources [10], [13]. Our two-source validation analysis, which used the depth-resolving intramuscular array as an independent measure of MU location, did not reveal a statistically significant systematic shift toward deeper MUs in the CRD. This suggests MU size, not location in terms of depth, is the primary factor for missed identification of MUs in initial decomposition. However, the validation sample contained relatively few matched MU pairs and was restricted to TA and VL, so this conclusion should be interpreted with caution.

Although the present analysis used convolutive blind-source-separation algorithms (CKC and FastICA variants) with fixed contrast functions, the core principle of CRD, which is removing dominant components to reveal previously masked sources, should generalize to other decomposition approaches. Any unsupervised or semi-supervised algorithm that identifies MUs in order of amplitude or variance will preferentially extract high-amplitude units first, leaving low-amplitude units in the residual. The approach is therefore compatible with alternative algorithms for offline MU identification including adaptive algorithms [12], MUAP-based methods [35], and deep-learning approaches [36]. However, the absolute yield gain will depend on the decomposition algorithm’s inherent ability to separate overlapping sources.

The principal practical disadvantage of the proposed pipeline is the additional manual editing effort required at each CRD iteration. In our experience, however, the net manual workload by including one CRD iteration does not increase by a factor of 2, although the number of automatic decomposition passes and manual editing processes is double. First, only MUs that pass strict initial identification quality criteria need to be retained from the original decomposition for the CRD approach to be useful. Ambiguous or noisy trains can be discarded rather than rescued through extensive editing, since their content is likely to be re-identified more cleanly in the next CRD iteration once the dominant MUs have been removed from the signal. Second, the residual signal contains fewer overlapping waveforms, resulting in relatively clean MU spike trains in CRD iterations. As a result, while still requiring some additional manual effort, we consider iterative CRD to be worth exploring for applications that would benefit from increased MU yield in ideal conditions. For easy integration of the approach into existing MU decomposition pipelines, we provide an open-source implementation of the residual computation and combination of CRD and original decomposition compatible with current state-of-the-art HD-sEMG decomposition implementations (see section Code Availability)

The present approach is limited to offline, post-hoc analysis and is not suitable for real-time or near-real-time MU identification due to the intermediate manual editing step. However, it demonstrates that the development of automated correction of identified spike trains is a promising future direction of MU decomposition research as the results prove the commonly observed errors and misidentifications in automated decompositions to be a bottleneck in increasing the number of observable motor units. Advances in that regard could increase the yield also in online, real-time decomposition scenarios. We further did not test the approach under dynamic contraction conditions, in pathological populations, or under fatigue and high-force nonstationary contractions. Finally, as with any surface EMG decomposition, a proportion of identified MUs may represent crosstalk from adjacent muscles, as MUs from adjacent muscles can be picked up by the recording electrodes and typically show smaller MUAP amplitudes due to the volume conduction of the muscle tissue [37]. However this potential confound is not specific to CRD but applies equally to the original decomposition and any decomposition approach.

## V. Conclusion

Our results establish Curated Residual Decomposition as a simple and reliable strategy for extending the motor unit yield of offline HD-sEMG decomposition in isometric conditions of healthy populations. The additional MUs identified through this approach are valid against an independent intramuscular reference, contribute meaningfully to previously unexplained signal power, and systematically expand the recorded MU pool toward physiologically informative smaller, lower-threshold units that are often undersampled in standard decomposition.

The consistency of results across three independent datasets spanning multiple muscles, different decomposition algorithm implementations, and various manual editing operators strongly supports the generalizability of the approach. Iterative CRD provides researchers and clinicians with a practical, optional enhancement step that can be readily integrated into existing HD-sEMG analysis workflows without requiring algorithmic modifications. The approach is straightforward to combine with future advances in decomposition algorithms, offering a flexible tool for improving motor unit identification completeness in offline neuromuscular assessment and motor control research. The development of automated spike train editing methods could extend the benefits of CRD to real-time or near-real-time applications, further expanding the utility of HD-sEMG decomposition techniques.

## Code Availability

An open-source implementation of the CRD pipeline in Python and Matlab compatible with the state-of-the-art decomposition implementations is available at https://github.com/NsquaredLab/curated_residual_decomposition_tool.

## Acknowledgment

M.O. and A.D.V. acknowledge the Deutsche Forschungsgemeinschaft (DFG, German Research Foundation) – Project DeMOTUS (523352235) and the European Research Council (ERC) – Project GRASPAGAIN (101118089).

## References

[1] E. Henneman, “Relation between size of neurons and their susceptibility to discharge,” Science (New York, N.Y.), vol. 126, no. 3287, pp. 1345–1347, 1957.

[2] D. Farina et al., “The extraction of neural strategies from the surface EMG,” Journal of Applied Physiology, vol. 96, no. 4, pp. 1486–1495, Apr. 2004.

[3] D. Farina et al., “The effective neural drive to muscles is the common synaptic input to motor neurons,” The Journal of Physiology, vol. 592, no. 16, pp. 3427–3441, 2014.

[4] R. M. Enoka and J. Duchateau, “Rate Coding and the Control of Muscle Force,” Cold Spring Harbor Perspectives in Medicine, vol. 7, no. 10, 2017.

[5] R. Merletti et al., “Analysis of motor units with high-density surface electromyography,” Journal of Electromyography and Kinesiology: Official Journal of the International Society of Electrophysiological Kinesiology, vol. 18, no. 6, pp. 879–890, Dec. 2008.

[6] A. Holobar and D. Zazula, “Multichannel Blind Source Separation Using Convolution Kernel Compensation,” IEEE Transactions on Signal Processing, vol. 55, no. 9, pp. 4487–4496, Sep. 2007.

[7] M. Chen and P. Zhou, “A Novel Framework Based on FastICA for High Density Surface EMG Decomposition,” IEEE Transactions on Neural Systems and Rehabilitation Engineering, vol. 24, no. 1, pp. 117–127, Jan. 2016.

[8] F. Negro et al., “Multi-channel intramuscular and surface EMG decomposition by convolutive blind source separation,” Journal of neural engineering, vol. 13, no. 2, Feb. 2016.

[9] S. Avrillon et al., “The identification of extensive samples of motor units in human muscles reveals diverse effects of neuromodulatory inputs on the rate coding,” eLife, vol. 13, p. RP97085, Dec. 2024.

[10] D. S. d. Oliveira et al., “Neural decoding from surface high-density EMG signals: influence of anatomy and synchronization on the number of identified motor units,” Journal of neural engineering, vol. 19, no. 4, Aug. 2022.

[11] A. Del Vecchio et al., “Tutorial: Analysis of motor unit discharge characteristics from high-density surface EMG signals,” Journal of Electromyography and Kinesiology, vol. 53, p. 102426, Aug. 2020.

[12] A. Grison et al., “Unlocking the full potential of high-density surface EMG: novel non-invasive high-yield motor unit decomposition,” The Journal of Physiology, vol. 603, no. 8, pp. 2281–2300, 2025.

[13] D. Farina et al., “A surface EMG generation model with multilayer cylindrical description of the volume conductor,” IEEE Transactions on Biomedical Engineering, vol. 51, no. 3, pp. 415–426, Mar. 2004.

[14] D. Farina et al., “Detecting the Unique Representation of Motor-Unit Action Potentials in the Surface Electromyogram,” Journal of Neurophysiology, vol. 100, no. 3, pp. 1223–1233, Sep. 2008.

[15] C. J. De Luca and P. Contessa, “Biomechanical benefits of the onion-skin motor unit control scheme,” Journal of Biomechanics, vol. 48, no. 2, pp. 195–203, Jan. 2015.

[16] M. Chen et al., “Automatic Implementation of Progressive FastICA Peel-Off for High Density Surface EMG Decomposition,” IEEE Transactions on Neural Systems and Rehabilitation Engineering, vol. 26, no. 1, pp. 144–152, Jan. 2018.

[17] C. Chen et al., “A peel-off convolution kernel compensation method for surface electromyography decomposition,” Biomedical Signal Processing and Control, vol. 85, p. 104897, Aug. 2023.

[18] S. Avrillon et al., “Tutorial on MUedit: An open-source software for identifying and analysing the discharge timing of motor units from electromyographic signals,” Journal of Electromyography and Kinesiology, vol. 77, p. 102886, Aug. 2024.

[19] M. Chen and P. Zhou, “Caution Is Necessary for Acceptance of Motor Units With Intermediate Matching in Surface EMG Decomposition,” Frontiers in Neuroscience, vol. 16, May 2022.

[20] S. Muceli et al., “Accurate and representative decoding of the neural drive to muscles in humans with multi-channel intramuscular thin-film electrodes,” The Journal of Physiology, vol. 593, no. 17, pp. 3789–3804, Sep. 2015.

[21] M. Osswald et al., “Task-Dependent Motor Unit Recruitment and Rate Coding Reveal Redistribution of Neural Drive in the Human Hand,” eLife, vol. 15, May 2026.

[22] M. Osswald et al., “Task-specific motor units in the extrinsic hand muscles control single-and multidigit tasks of the human hand,” Journal of Applied Physiology, vol. 138, no. 5, pp. 1187–1200, May 2025.

[23] S. Avrillon, “Data for the preprint “The decoding of extensive samples of motor units in human muscles reveals the rate coding of entire motoneuron pools”,” 2023, dataset; licensed under CC BY 4.0.

[24] F. Hug et al., “Analysis of motor unit spike trains estimated from high-density surface electromyography is highly reliable across operators,” Journal of Electromyography and Kinesiology, vol. 58, p. 102548, Jun. 2021.

[25] F. Hug et al., “Analysis of motor unit spike trains estimated from high-density surface electromyography is highly reliable across operators,” 2021, dataset; licensed under CC BY 4.0.

[26] A. Holobar et al., “Experimental Analysis of Accuracy in the Identification of Motor Unit Spike Trains From High-Density Surface EMG,” IEEE Transactions on Neural Systems and Rehabilitation Engineering, vol. 18, no. 3, pp. 221–229, Jun. 2010.

[27] H. R. Marateb et al., “Accuracy assessment of CKC high-density surface EMG decomposition in biceps femoris muscle,” Journal of Neural Engineering, vol. 8, no. 6, p. 066002, Oct. 2011.

[28] M. Chen et al., “Two-Source Validation of Progressive FastICA Peel-Off for Automatic Surface EMG Decomposition in Human First Dorsal Interosseous Muscle,” International Journal of Neural Systems, vol. 28, no. 09, p. 1850019, Nov. 2018.

[29] D. Farina and A. Holobar, “Characterization of Human Motor Units from Surface EMG Decomposition,” Proceedings of the IEEE, vol. 104, no. 2, pp. 353–373, Feb. 2016.

[30] F. Negro and D. Farina, “Factors Influencing the Estimates of Correlation between Motor Unit Activities in Humans,” PLoS ONE, vol. 7, no. 9, p. e44894, Sep. 2012.

[31] X. Hu et al., “Control of motor unit firing during step-like increases in voluntary force,” Frontiers in Human Neuroscience, vol. 8, Sep. 2014.

[32] C. J. De Luca and Z. Erim, “Common drive of motor units in regulation of muscle force,” Trends in Neurosciences, vol. 17, no. 7, pp. 299–305, Jan. 1994.

[33] G. Valli et al., “Lower limb suspension induces threshold-specific alterations of motor units properties that are reversed by active recovery,” Journal of Sport and Health Science, vol. 13, no. 2, pp. 264–276, Mar. 2024.

[34] J. Duchateau and K. Hainaut, “Effects of immobilization on contractile properties, recruitment and firing rates of human motor units.” The Journal of Physiology, vol. 422, no. 1, 1990.

[35] C. Chen et al., “A motor unit action potential-based method for surface electromyography decomposition,” Journal of NeuroEngineering and Rehabilitation, vol. 22, no. 1, p. 60, Mar. 2025.

[36] A. K. Clarke et al., “Deep Learning for Robust Decomposition of High-Density Surface EMG Signals,” IEEE Transactions on Biomedical Engineering, vol. 68, no. 2, pp. 526–534, Feb. 2021.

[37] C. M. Germer et al., “Surface EMG cross talk quantified at the motor unit population level for muscles of the hand, thigh, and calf,” Journal of applied physiology (Bethesda, Md. : 1985), vol. 131, no. 2, pp. 808–820, Aug. 2021.

